# Lateral inhibition governs ancestral cellular patterning in fossil and extant liverworts

**DOI:** 10.64898/2025.12.27.696693

**Authors:** Josep Mercadal, Susan Tremblay, Leonie Kraska, Martin A. Hutten, Pau Formosa-Jordan

## Abstract

The incompleteness of the fossil record and the absence of genetic information limit our ability to understand ancient developmental processes and reconstruct their evolution. Here, we use comparative and mathematical analyses to uncover an evolutionarily conserved patterning mechanism underlying liverwort oil body cell specification. We demonstrate that patterns of oil body cell homologs in fossils of the oldest known liverwort (*Metzgeriothallus sharonae*) exhibit the spatial organization characteristic of lateral inhibition. A mechanism with local activation and lateral inhibition not only reproduces the patterns observed in fossils, but also oil body cell patterns in the extant liverworts *Treubia lacunosa, Apotreubia nana*, and *Marchantia polymorpha*. Our results point to an ancestral patterning mechanism that has been modulated by lineage-specific selective pressures over evolutionary time.

---

Evolutionary developmental biology has historically relied on comparative anatomy, morphometrics, and systematics, with fossil evidence playing a crucial role in inferring developmental patterns and their evolutionary relationships^1–3^. While modern approaches to understanding the evolution of development have gravitated towards molecular techniques, these remain inadequate when confronted with ancient fossil data, where genetic material is rarely preserved. In particular, reconstructing dynamical aspects of development—such as morphogenesis and cell-specification mechanisms—remains challenging and often underexplored. Recent advances in imaging and mathematical modeling have revived comparative approaches^4–8^, making it possible to recognize morphological fingerprints of developmental mechanisms in living and extinct organisms^9–13^. In animals, these include the evolution of teeth development^14–17^, the fin-to-limb transition^18–20^, and patterning of epidermal structures^21–23^, among others. While less studied in this context, plants offer a unique opportunity to investigate ancestral developmental mechanisms, as preserved cell walls and fixed cell positions facilitate the recovery of developmentally informative anatomy^24–28^. Evidence includes the presence of ancient polar auxin transport from growth patterns in wood and vascular tissues^29,30^, or polyploidy and conserved patterning in fossil stomata^31–33^.

A striking example of well-preserved cellular patterning can be observed in the Middle Devonian (*c*. 388 Ma) fossil *Metzgeriothallus sharonae*^34^, the oldest known liverwort. A defining feature of these fossils is the presence of dark, scattered cells that closely resemble oil body (OB) cells of modern liverworts (Fig. 1A), suggesting evolutionary homology^35,36^ and implicating these structures in the chemical defense of liverworts against arthropod herbivory^37^. Dark cells are distributed more or less evenly throughout the thallus wings, often forming characteristic *salt-and-pepper* patterns (Fig. 1). This spatial arrangement is associated with the phenomenon of lateral inhibition, where adjacent cells compete to differentiate into distinct types by inhibiting the same fate as their neighbors, generating regularly spaced patterns^38–40^.

**Figure 1:**
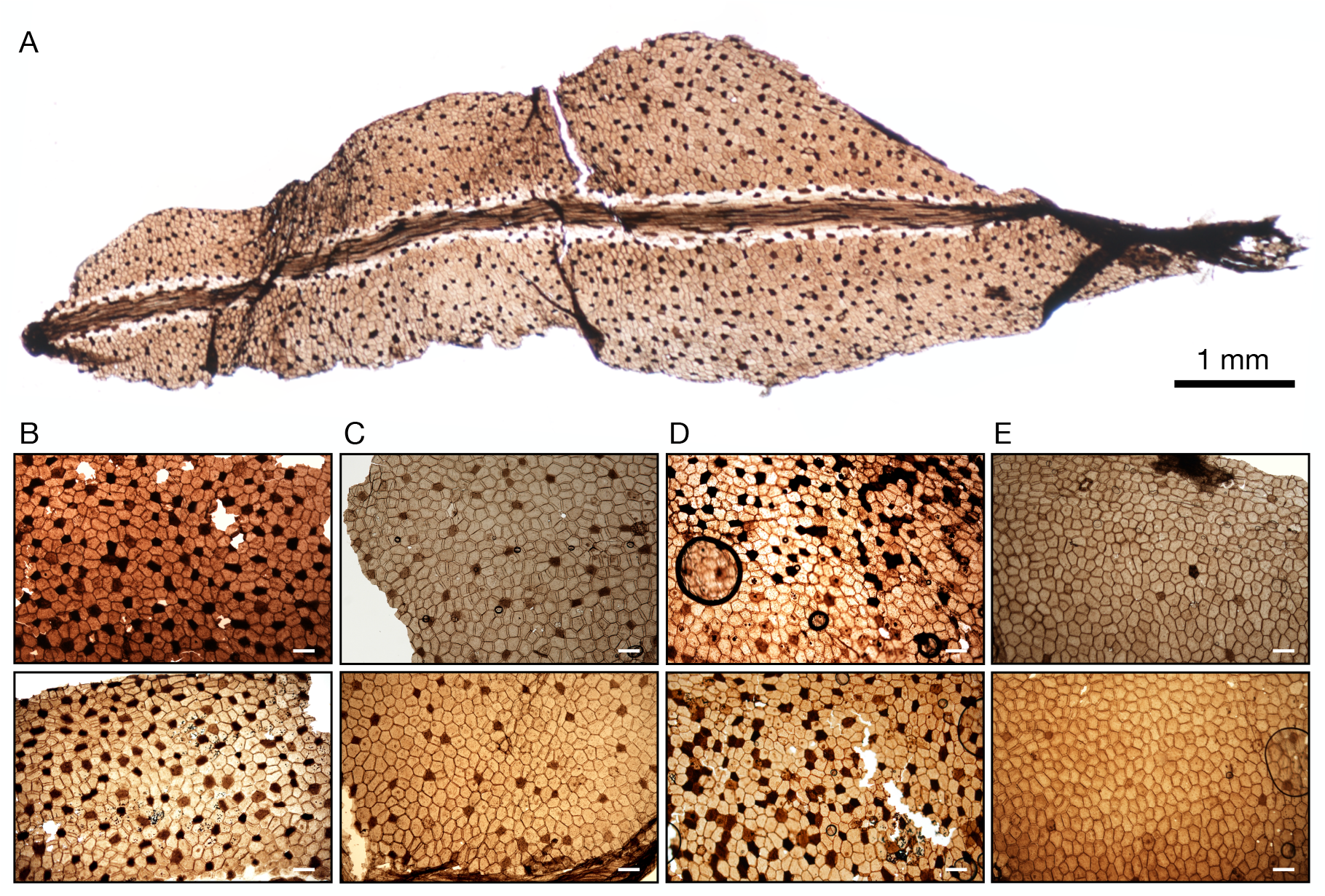
Light micrographs of the Middle Devonian fossil liverwort *Metzgeriothalus sharonae*. **A**. Thallus comprised of midrib and unistratose wings with scattered dark cells. Scale bar = 1 mm. **B–E**. Representative sections of *M. sharonae* showing sections of different thalli, where scattered dark cells can be readily discerned. These dark cells are regularly distributed in space, mostly isolated (idioblasts) or forming small clusters of two to three cells (B, C). Occasionally, larger, mostly linear clusters of dark cells can also be observed (D). In a few sections, dark cells appear only occasionally and in idioblastic form (E). Scale bars = 0.1 mm.

Here, we show that dark cell patterns in *M. sharonae* exhibit the characteristic hallmarks of lateral inhibition, including an overrepresentation of isolated dark cells (idioblasts) and a distinctive spatial regularity. We demonstrate that the spatial statistics of dark cells are consistent with cell specification mediated by local activation and lateral inhibition, accounting for the full diversity of patterns observed in fossil samples. Finally, we show that the same cell specification mechanism can reproduce the spatial organization of OB cells in the extant liverworts *Treubia lacunosa, Apotreubia nana*, and *Marchantia polymorpha*, supporting the evolutionary conservation of an ancestral patterning process. Our framework explains lineage-specific phenotypic transitions—such as the complete loss of OBs or the shift from ubiquitous to idioblastic OB distribution—as outcomes of a conserved patterning module being repurposed over evolutionary time.

## Dark cell pattern diversity and spatial statistics

We collected *M. sharonae* material from the Cairo quarry in the Catskill Delta deposit of eastern New York state, well-known for its diverse macroflora, and the only known liverwort-dominated community from the Paleozoic^34,36,37^. From a total of 120 collected *M. sharonae* samples, we selected 58 sections based on the size (> 1 mm^2^), good preservation of cellular structures, and clearly defined dark cells. The total number of cells in each section ranged from ≈ 400 to 900. Due to the variability in frequency and regularity of dark cell patterns, we classified sections into four categories: I) regular patterns with high dark cell frequency (*ρ*_*I*_ ≥ 0.19), where dark cells form salt-and-pepper patterns with a separation of 1 to 2 epidermal cells (Fig. 1B); II) regular patterns with low dark cell frequency (*ρ*_*II*_ < 0.19), where dark cells are separated by 2 to 5 epidermal cells (Fig. 1C); III) mixed patterns, where the regularity of dark cell patterns is not obvious due to the presence of gaps and large clusters (Fig. 1D); and IV) patterns with very few scattered dark cells (Fig. 1E). Most sections belong to category II (*n*_*II*_ = 43), whereas only a few belong to categories I (*n*_*I*_ = 7), III (*n*_*III*_ = 5), and IV (*n*_*IV*_ = 3). Because dark cells are rare in type IV sections, we excluded them from the statistical analyses, leaving a total of *n* = 55 fossil sections.

We used the following metrics to characterize the spatial distribution of dark cells (Supplementary Materials; Figs. S1 and S2): dark cell frequency *ρ*, i.e., the proportion of dark cells relative to the total number of cells in the tissue; the distribution of sizes of dark cell clusters *f*_*k*_, where *k* represents the cluster size; the distribution of cell distances between pairs of neighboring dark cells *D*_*j*_, where *j* stands for the number of separating epidermal cells; and the spatial autocorrelation via Moran’s *I* statistic. We compared these metrics with those obtained with a random model of cell specification, in which each cell becomes dark with probability *p* or epidermal with probability 1 − *p* (Fig. 2; Fig. S3). We performed simulations of the random model on irregular cell lattices with cell numbers similar to those in fossil samples (Fig. 2A). In fossils, dark cells appear with a frequency *ρ* = 0.16 ± 0.2 (Fig. 2A,B). The distribution of cluster sizes shows a strong excess of idioblasts (clusters of size *k* = 1), significantly more than what would be expected by random specification (Fig. 2D). In contrast, larger clusters occur at considerably lower proportions than expected by chance (Fig. 2D), and typically take the form of linear arrays, which are distinctive hallmarks of lateral inhibition^41,42^. Most dark cells are separated by 1 to 3 cells, with an average of ≈ 2.7 cells (Fig. 2E). This regularity contrasts with the random model, where dark cell separation is significantly more variable (Fig. 2E; Fig. S2).

**Figure 2:**
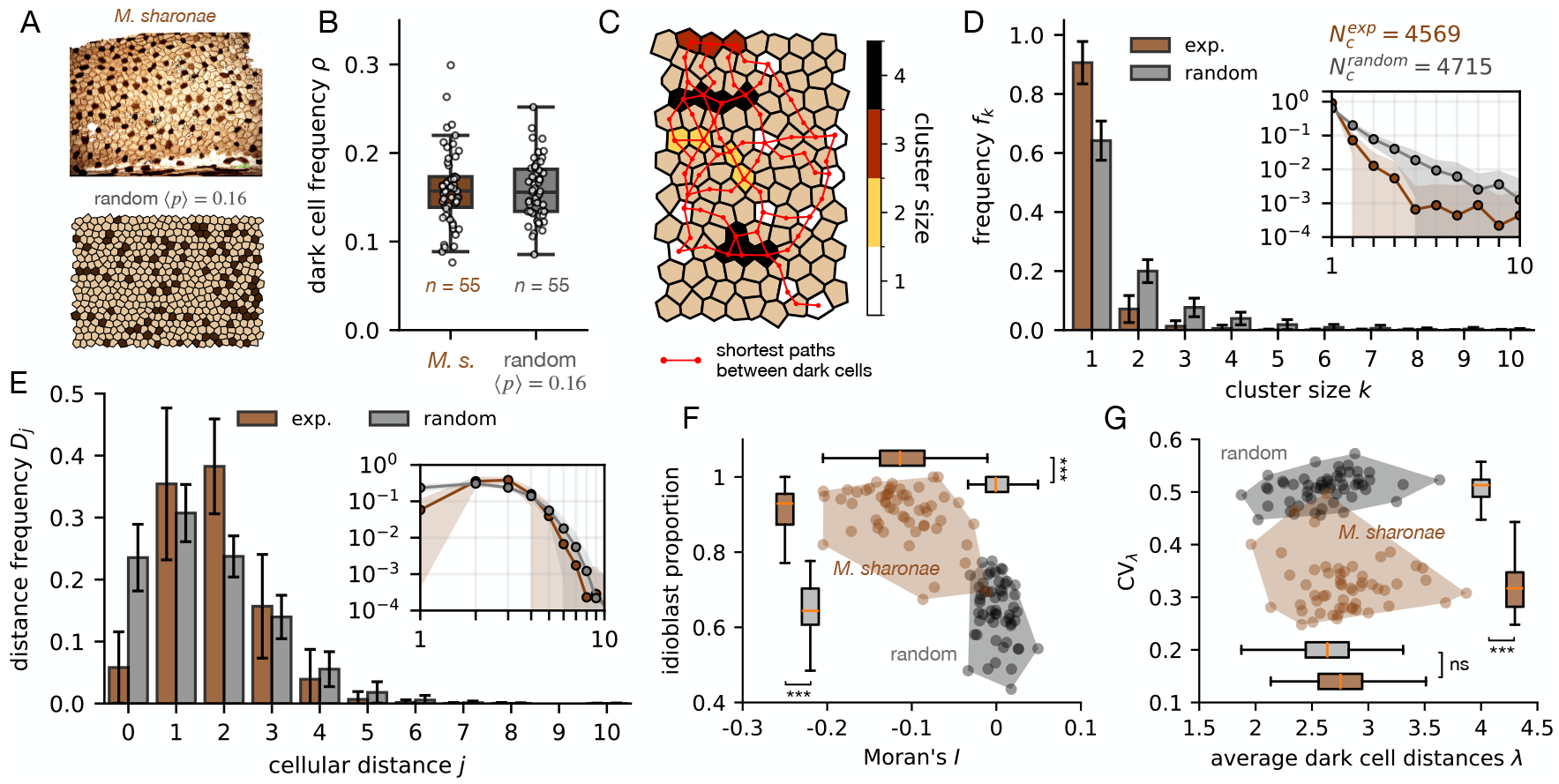
Non-randomness of dark cell patterns in *M. sharonae* fossils. **A**. Representative section of *M. sharonae* showing a dark cell pattern (top) and a simulated random pattern with similar dark cell frequency (bottom). **B**. Boxplots showing the frequency of dark cells for fossil sections (brown, *n* = 55) and for random patterns (gray, *n* = 55) with dark cell probability *p* = 0.16 × (1 + 0.7*𝒰* [−0.5, 0.5]), where *𝒰* [−0.5, 0.5] denotes a uniform distribution of random numbers between −0.5 and 0.5. With this choice, random patterns exhibit a frequency distribution similar to that of fossil samples. **C**. Metrics used to determine the spatial statistics of dark cell patterns, including the cluster sizes in different colors and the shortest paths between neighboring dark cells in red (see Fig. S1 and Supplementary Materials for a complete description of these metrics). **D**. Distribution of cluster sizes in fossil patterns (brown) compared to random patterns (gray). The inset shows the same data on a log-log scale. *N*_*c*_ stands for the total number of clusters analyzed. **E**. Distribution of distances (shortest paths) between dark cells in fossils (brown) and random patterns (gray). **F**. Morphospace showing the proportion of idioblastic dark cells versus Moran’s *I* in fossil (brown) and random (gray) patterns. **G**. Morphospace showing the average distances between dark cells versus their corresponding coefficient of variation of fossil (brown) and random (gray) patterns. In panels B, F, and G, box widths represent interquartile ranges (IQR = Q3-Q1), horizontal lines represent medians, and whiskers extend from Q1-1.5IQR to Q3+1.5IQR. Error bars and shaded areas in D and E denote ± standard deviations. The shaded regions in panels F and G represent the convex hulls of each dataset. Statistical significance was assessed using the Mann-Whitney U-test; *** *p* < 0.01; ns: non-significant.

To better understand how dark cell patterns compare to a random model of cell specification, we located each fossil sample in the morphospaces of Moran’s *I* statistic versus idioblast proportion, and average dark cell distance ⟨*λ*⟩ versus *CV*_*λ*_ —the coefficient of variation of these distances (see Supplementary Materials for how these metrics are defined). Our results show that in these spaces, dark cell patterns cluster without overlapping with random patterns (Fig. 2F,G). Dark cell patterns have a high proportion of idioblasts and negative values of Moran’s *I*, revealing a spatial organization that the random model cannot reproduce (Fig. 2F). While the average distance between dark cells is similar between the fossils and the random model, the *CV*_*λ*_ is significantly lower in the fossils (Fig. 2G), reflecting lower variability in these distances and a characteristic pattern regularity. Altogether, these results show that the statistical properties of dark cell patterns in fossils are incompatible with random cell specification. This suggests that the mechanism of dark cell specification in *M. sharonae* was not random—as would be expected by a cell-autonomous process—but likely involved cell-to-cell interactions via lateral inhibition, preventing adjacent dark cells from adopting the same fate and generating the observed spatial regularity.

## A model with local activation and lateral inhibition reproduces the diversity of dark cell patterns

Developmental mechanisms generating cell-type-specific patterning typically rely on cell-to-cell communication through variants of lateral inhibition^43–45^. While the genetic and molecular factors involved in patterning might differ significantly across biological systems, the dynamics and spatial statistics are often telltale signs of the underlying developmental processes and offer crucial information for distinguishing between mechanisms^46,47^. Due to the lack of genetic and dynamical information, we compared the spatial statistics of dark cell patterns in *M. sharonae* with those obtained with a minimal cellular model of pattern formation, where a local activator *u* induces the activity of a mobile inhibitor *v*^42,48^ (Fig. 3A; Supplementary Materials). At each time point, the state of each cell is described by the values of *u*(*t*) and *v*(*t*). We assume the activator is constantly produced, self-activates, and is degraded linearly and by the inhibitor (Fig. 3B). The inhibitor is also constantly produced, is induced by the activator, is linearly degraded, and can diffuse to adjacent cells (Fig. 3B). We explored the steady-state patterning properties of the model in an irregular cellular tissue by varying the parameters *γ*—the maximum rate at which *u* activates *v*—and *α*_0_—the basal production rate of *u* (Fig. 3C; Fig. S4). We use the stationary values of *u* as a proxy for cell fate, with high *u* indicating dark cells and low *u* indicating epidermal cells.

**Figure 3:**
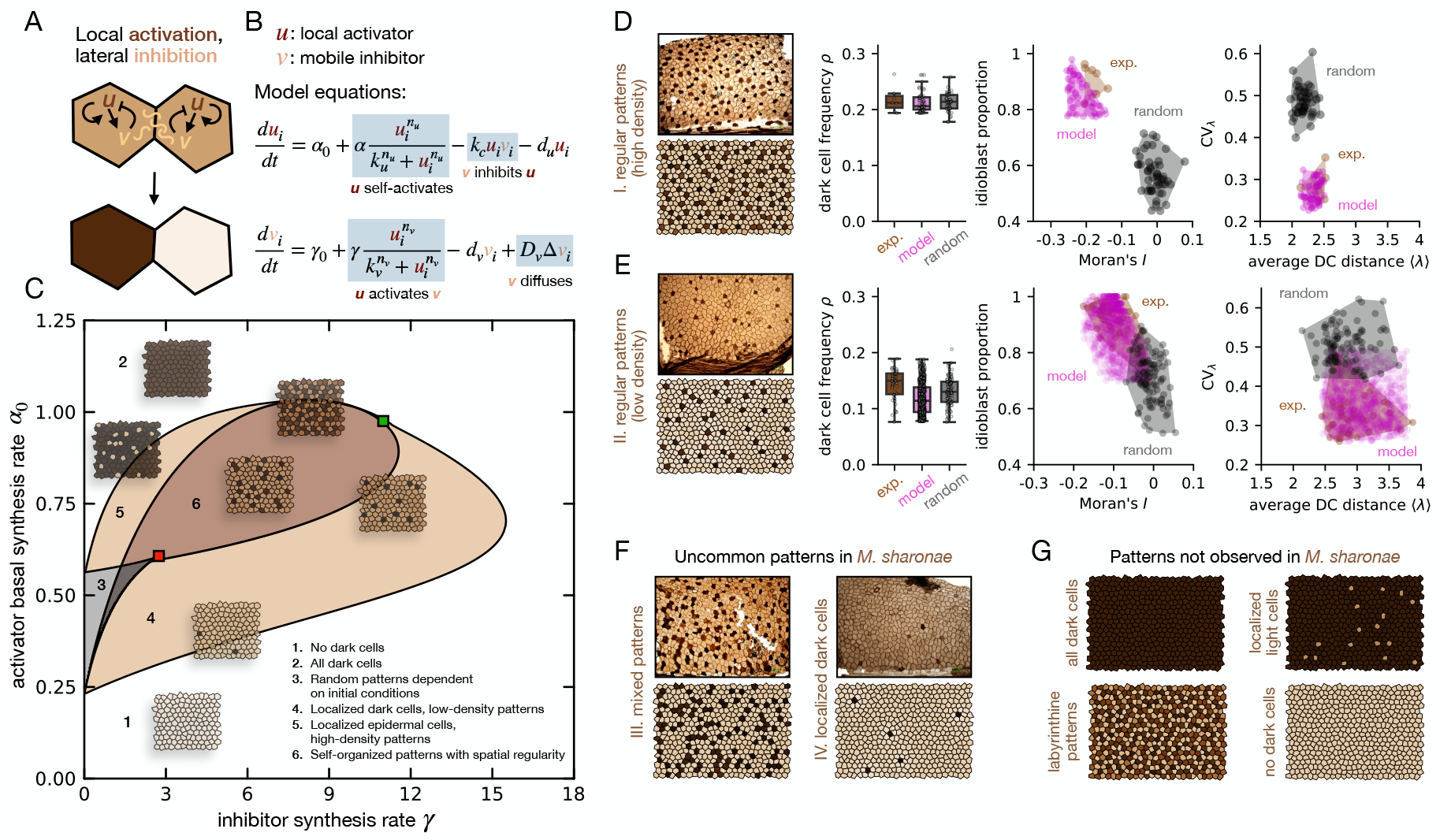
A model with local activation and lateral inhibition reproduces the diversity of dark cell patterns in *M. sharonae* fossils. **A**. Sketch of the interactions in a cellular model of local activation and lateral inhibition. The state of two adjacent undifferentiated cells can be destabilized due to diffusive lateral inhibition, priming them to adopt opposite fates. **B**. Equations of the mathematical model corresponding to the interactions depicted in A. The key regulations are highlighted in blue. **C**. Stability diagram showing different patterning regimes obtained with the mathematical model as a function of the parameters *γ* and *α*_0_ (Supplementary Materials; Fig. S4). The red square denotes a cusp point where two saddle-node bifurcations collide and annihilate; the green square denotes the collision between a subcritical Turing bifurcation and two saddle-node bifurcations. Snapshots of representative simulated patterns are displayed across the stability diagram. **D**. Comparison of the spatial statistics of high-density fossil samples (brown; *n* = 7) with the mechanistic (pink; *n* = 55) and random (gray; *n* = 100) models (see Fig. S5). **E**. Comparison of low-density fossil samples (brown; *n* = 43) with the mechanistic (pink; *n* = 826) and random (gray; *n* = 100) models (see Fig. S5). **F**. In a few sections, dark cells are inhomogeneously distributed (left) or appear only occasionally (right). Statistics were not performed due to the small number of sections of these types. Nevertheless, the mathematical model can qualitatively reproduce the corresponding patterns (Fig. S5). **G**. Several patterns predicted by the model are not observed in the fossils. Cell colors represent the values of the variable *u* in each cell, ranging from *u* ∈ [0.2, 1.2] (see Fig. S4). The values of *α*_0_ and *γ* used for panels D–G are specified in Fig. S5. The rest of the parameters are detailed in Table S1.

Analyzing the model through bifurcation theory and linear stability analysis^38,49^ reveals different patterning regimes (Fig. 3C; Fig. S4). If *α*_0_ is small, the activator is not strong enough to induce cell differentiation, and no dark cells emerge (Fig. 3C; region 1). Conversely, for large *α*_0_, all cells become dark (region 2). Patterns with different dark cell frequencies and regularities can emerge for intermediate values of (*γ, α*_0_). The system has a bistable region where cellular fates are strongly influenced by initial conditions (region 3). Here, lateral inhibition does not play a significant role because all possible cell combinations are stable (Fig. S4). The model can also generate localized patterns with different densities (regions 4 and 5) that are triggered only by specific initial conditions (Fig. S4). Finally, for a wide range of (*γ, α*_0_), the system can generate self-organized patterns (region 6), whereby ordered spatial patterns emerge spontaneously due to the amplification of inhomogeneous perturbations. Patterns across this region differ in their spatial statistics, including dark cell frequency, idioblast proportion, and average dark cell distance (Fig. S4). Mixed patterns—where periodic domains coexist with more random-looking patterns—can be obtained when the values of (*γ, α*_0_) are close to the bistable region (regions 4,5,6 close to region 3; Fig. S4).

We performed a parameter sweep over the pairs (*γ, α*_0_) to test whether the model could reproduce the statistical metrics of fossil patterns (dark cell frequency, idioblast proportion, Moran’s *I*, average dark cell distance ⟨*λ*⟩, and *CV*_*λ*_) across the four categories of our classification (Fig. 3D,F; Fig. S5). For each category, we identified the pairs (*γ, α*_0_) that generated patterns in which all metrics simultaneously fell within their observed experimental ranges (Supplementary Materials; Fig. S5). We found that the model could reproduce all the observed phenotypes, generating patterns in morphospace regions that virtually overlap or contain the experimental results (Fig. 3D,E). Notably, the model yields multiple parameter pairs that satisfy all these conditions (Fig. S5), underscoring the mechanisms’ ability to generate similar phenotypes through different routes. Uncommon patterns, such as those with no well-defined regularity or very few localized dark cells, could also be reproduced by the model (Fig. 3F; Fig. S5). Additionally, the model can generate phenotypes that are not observed in fossil samples. These include tissues completely covered in dark cells, tissues with mostly dark cells and few localized light cells, labyrinthine patterns, and tissues without dark cells (Fig. 3G; Fig. S4,S5).

Taken together, these results demonstrate that a cell specification mechanism through local activation and lateral inhibition can explain the full variation of dark cell patterns in *M. sharonae*. This flexibility in generating diverse patterning phenotypes highlights the evolutionary potential of the mechanism, enabling adaptive responses to varying selective pressures.

## An evolutionarily conserved patterning mechanism for oil body cell specification

Our results suggest a cell patterning mechanism underlying the specification of dark cells in *M. sharonae*, the oldest known fossil liverwort. Dark cells in *M. sharonae* and other Paleozoic liverworts are considered homologous to the OB cells of modern liverworts^36,37^. Hence, to investigate whether this mechanism has been conserved across the liverwort lineage, we analyzed the spatial distribution of OB cells in thalli of the extant liverworts *Treubia lacunosa, Apotreubia nana*, and *Marchantia polymorpha* (Fig. 4). In *T. lacunosa* and *A. nana*, cells harboring OBs are on average larger than non-OB cells, and most appear as idioblasts (Fig. 4A,C). Notably, they occur at a similar frequency as in *M. sharonae* (*ρ* ≈ 0.15; Fig. 4B,D). Most of the OB cell patterns in *T. lacunosa* and *A. nana* are inconsistent with random cell specification, despite some overlap in the regions occupied by the morphospaces (Fig. 4B,D). Instead, both the idioblast proportion, Moran’s *I*, average OB cell distance ⟨*λ*⟩, and its coefficient of variation *CV*_*λ*_ are consistent with patterns shaped by lateral inhibition. To test whether our model could reproduce the observed statistics, we looked for pairs (*γ, α*_0_) whose statistical metrics reproduced OB cell patterns in *T. lacunosa* and *A. nana*. We found several pairs that fulfilled all the conditions (Fig. 4B,D, pink; Fig. S6), spanning a morphospace that virtually overlapped or contained the experimental results. Therefore, OB cell patterns in *T. lacunosa* and *A. nana* are consistent with cell specification governed by local activation and lateral inhibition.

**Figure 4:**
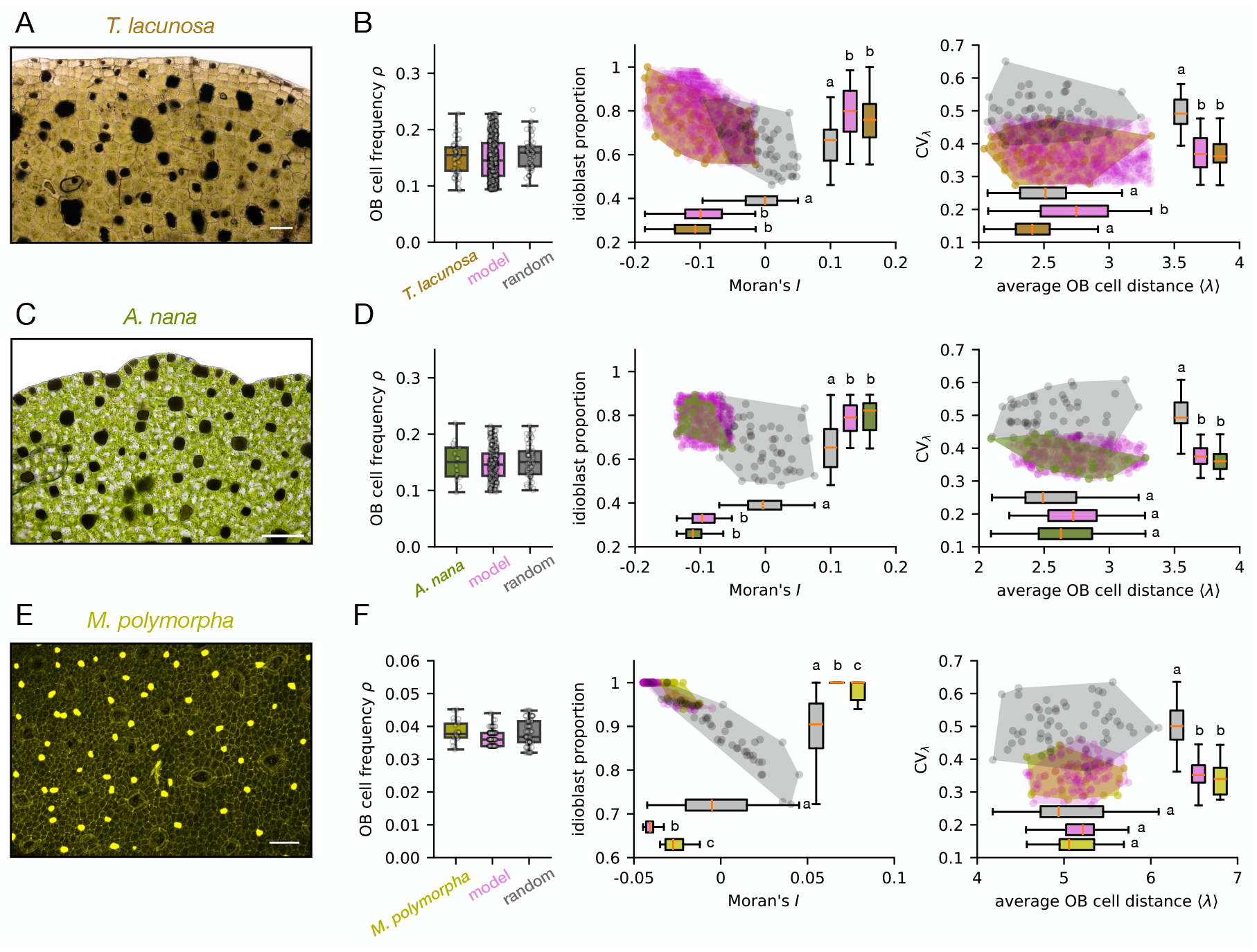
Lateral inhibition patterning of oil body cells is conserved across liverwort taxa. **A**. Distribution of OB cells in *T. lacunosa*. **B**. Spatial statistics comparing experimental OB cell patterns in *T. lacunosa* (brown; *n* = 50) with patterns obtained with the mechanistic (pink; *n* = 777) and random (gray; *n* = 55) models. **C**. Distribution of OB cells in *A. nana*. **D**. Spatial statistics comparing OB cell patterns in *A. nana* (green; *n* = 16) with patterns obtained with the mechanistic (pink; *n* = 305) and random (gray; *n* = 55) models. **E**. *M. polymorpha* thallus stained with BODIPY, exposing oil bodies and cell boundaries ( Fig. S7). **F**. Spatial statistics comparing OB cell patterns in *M. polymorpha* (gold; *n* = 21) with patterns obtained with the mechanistic (pink; *n* = 84) and random (gray; *n* = 55) models. Statistical differences in panels B, D, and F were tested using ANOVA followed by Tukey HSD (*p* < 0.01); letters indicate statistically significant groups. Scale bars in A, C, and E: 0.1 mm.

Finally, we tested whether our model could reproduce OB cell patterns in *M. polymorpha*. In *M. polymorpha* thalli, OB cell frequency is lower than in fossils, *T. lacunosa* and *A. nana*, and almost exclusively in idioblastic form (Fig. 4E,F; Fig. S8). Due to this lower frequency, the average distance between OB cells is higher than in the other species. By contrast, the coefficient of variation spans the same scale, suggesting that despite differences in frequency and OB cell distance, the pattern variability is conserved. Comparing these distributions with what a random model with the same OB cell frequency reveals that patterns in *M. polymorpha* are also shaped by lateral inhibition. Indeed, our model reproduced all the spatial statistics associated with these patterns (Fig. 4F; Fig. S6).

Taken together, our results show that a mechanism with local activation and lateral inhibition can reproduce the spatial patterns of OB cells not only in *M. sharonae* fossils, but also in the extant liverworts *T. lacunosa, A. nana*, and *M. polymorpha*. This consistency across distantly related liverwort taxa points to a deeply conserved developmental mechanism underlying liverwort OB cell specification.

## Discussion

Oil bodies play important roles in lipid and secondary metabolite storage, thereby contributing significantly to plant defense mechanisms^50–54^. The regular and idioblastic nature of OB cells might reflect a strategy to minimize unnecessary overlap, potentially optimizing their protective or signaling functions. Despite their putative defensive role, the frequency and spatial distribution of OB vary among liverwort taxa. In Haplomitriopsida, taxa in the genus *Haplomitrium* have several OBs in all cells. In contrast, those in the *Treubiales* (like *T. lacunosa* and *A. nana*) have a single OB in idioblastic cells^52,55^ (Fig. 5A). In Marchantiopsida, most taxa have a single OB in idioblastic cells, while in Jungermanniopsida (where *M. sharonae* is currently assigned^34,36^), most taxa have OBs in all cells^52,55^ (Fig. 5A). The three liverwort classes are estimated to have diverged from a common ancestor no later than the Late Silurian, and last shared a common ancestor with mosses in the Late Ordovician^56,57^. These divergence times highlight the ancient existence of OBs as a crucial adaptation to life on land.

**Figure 5:**
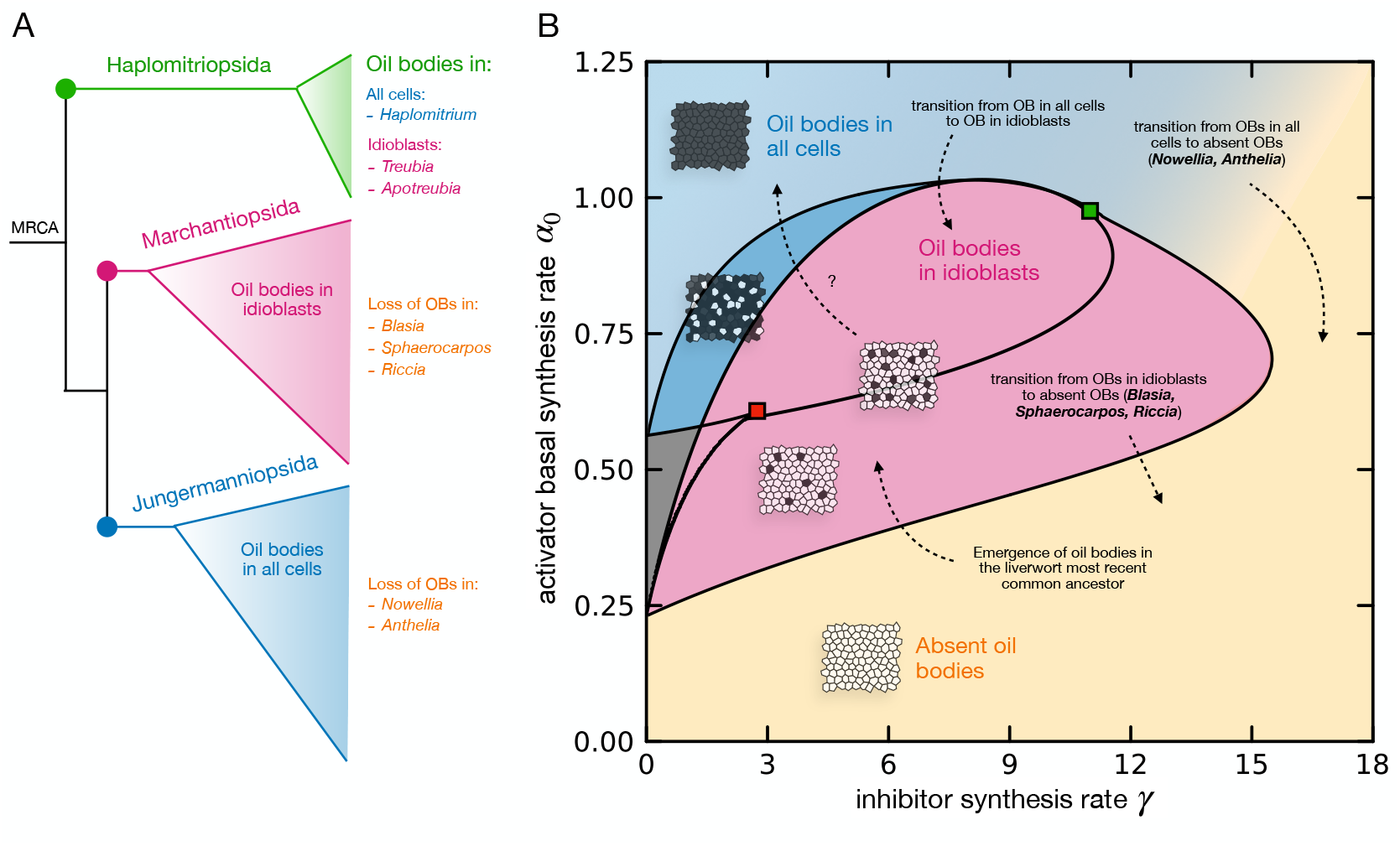
Oil body cell evolution through the lens of a flexible patterning module. **A**. Oil body cell distribution across the liverwort phylogeny. Branch colors indicate the general spatial arrangement of OB cells in the corresponding lineages: Haplomitriopsida (green), Marchantiopsida (magenta), and Jungermanniopsida (blue), together with a description of how OB cells are distributed. MRCA: most recent common ancestor. **B**. Patterning regimes (Fig. 3C) showing where OBs appear in all cells (light blue), in idioblastic cells (magenta), or are absent (yellow). Phenotypic transitions during evolution can be interpreted as excursions in this space. For instance, the transition from having OBs in all cells to their disappearance (as seen in *Nowellia* or *Anthelia*) can be interpreted either as a smooth change (e.g, through slowly changing environments) or as a sharp transition (e.g., through strong mutations). Similarly, the transition from OBs in idioblasts to OB loss (as occurs in many genera of *Marchantiopsida*) can be interpreted as the crossing of a bifurcation boundary.

The variation in type and spatial organization of OBs suggests a developmental flexibility that can be co-opted to produce more specialized adaptations suited to changing ecological contexts. Notably, most of the phenotypic variation in OB cell patterns was already present in *M. sharonae*, suggesting that developmental plasticity could have supported timely responses to the changing environments and emerging biotic stresses of Devonian landscapes. In our modeling framework, phenotypic variation reflects the mechanisms’ ability to explore different regions of the patterning space (Figs. 3C and 5B). Modulating the relative effect of the activator and inhibitor constitutes a route for evolutionary change by shifting the system between different stable states^4,7^. These adjustments can lead to bifurcations in the developmental dynamics, where small variations in the parameters result in large phenotypic changes. In this regard, phenotypic transitions such as the loss of OBs or the shift from ubiquitous to idioblastic OB distribution reflect incursions into qualitatively distinct patterning domains (Fig. 5B), which can occur as a result of, e.g., genetic mutations, biotic stresses, or environmental perturbations. We thus hypothesize that the patterning mechanism underlying OB cell specification arose early in liverwort evolution and has persisted despite extensive diversification, being repurposed over time due to lineage-specific selective pressures.

Despite being one of the defining synapomorphies of the group, OBs have rarely been used to explore evolutionary relationships within the liverwort clade, except for a few studies^35,58^. This is striking considering that the internal and spatial variability of OB cells is taxonomically informative and could offer insights into the adaptive strategies of early land plants. Idioblastic dark cells, very similar in structure and distribution to those of *M. sharonae*, have also been described in fossils from the early liverworts *Pallaviciniites devonicus*^36,59^ and *Treubiites kidstonii*^60,61^, and in Paleozoic mosses^62–64^, suggesting OBs may have originated much earlier than currently assumed and could even predate the divergence of bryophyte lineages.

Recent studies in *M. polymorpha* are shedding light on the molecular and developmental mechanisms underlying OB cell specification. The transcription factors MpC1HDZ, MpERF13, and MYB02 act as activators of OB cell specification, as loss-of-function mutants lack OB cells altogether^53,65–67^. Notably, MpC1HDZ and MpERF13 activate the expression of MYB02^67^. The transcription factor MpTGA represses the expression of both MpC1HDZ and MpERF13, and loss-of-function mutants of MpTGA show a dramatic increase in OB cells. Our model can qualitatively reproduce all these phenotypes (Fig. S9). In addition to modulating the spatial distribution of OB cells, several of these genes are necessary for the synthesis of OB-specific compounds. TGA represses MpSYP12B, an OB-resident SNARE protein that localizes in the OB membrane^53,68^, and upregulates terpenoid biosynthesis enzymes encoded by TPS and MTPSL^68^. The transporter MpABCG1 was recently shown to mediate the OB accumulation of sesquiterpenes, compounds that contribute to herbivore resistance^69^. Finally, the immune regulator MpNPR has been shown to control the expression of MpERF13, MpSYP12B, MpMYB02, and MpABCG1, thus linking the defensive function of OBs with the developmental processes underlying OB cell distribution. Despite the lack of any feedback mechanism that could function as an activator-inhibitor system capable of cell-type patterning, these studies are beginning to link the components of what appears to be a complex gene regulatory network (Fig. S10), providing a foundation for future reconstructions of the developmental mechanisms underlying OB formation and evolution.

## Supporting information

Supplementary Materials (including Materials and Methods, Supplementary Figures and Supplementary Tables)

## ACKNOWLEDGEMENTS

J.M. thanks the members of the Formosa-Jordan group for valuable discussions. S.T. thanks Taehee Han and Victoria Chu for providing laboratory assistance for the selection and preparation of fossils, David Glenny and Bill Malcolm for field support in New Zealand, and Linda Hernick for guidance and advice during field work in New York. L.K. thanks Arezki Boudaoud for hosting and support, and Pierre Mahou for providing access to the microscope facilities and technical support. M. H. thanks Edward Kulack for help in preparing LIDAR data and James Walton for help with fieldwork.

## Author contributions

J.M. designed the research, performed the image analysis, and analyzed the mathematical models. S.T. collected the samples of *M. sharonae* and *T. lacunosa*, and generated the images. L.K. produced the confocal images of *M. polymorpha*. M.H. collected the samples of *A. nana* and generated the images. J.M. wrote the manuscript with input from the other authors.

## Data and materials availability

All data reported in this paper will be shared by the lead contact upon request. All original code for data analysis and simulations is available in https://github.com/josepmercadal/fossils.

