## Supplementary Materials (including Materials and Methods, Supplementary Figures and Supplementary Tables) for "Lateral inhibition governs ancestral cellular patterning in fossil and extant liverworts"

#### Sample collection and processing

##### *Metzgeriothallus sharonae*

*Metzgeriothallus sharonae* material originates from the Cairo quarry, New York State, located south of New York State Route 145 at the Cairo Highway department headquarters, 3.22 km NW of Cairo, Greene County, New York (42.32ºN and 74.04ºW, NAD 83). The material was collected from three locations approximately 3m apart along a 25m-wide exposed lens of dark gray shales and mudstones. Shale fragments were immersed in 48% hydrofluoric acid to dissolve the shale matrix. The isolated carbonaceous fossils were rinsed in water until a neutral pH was obtained, and then they were rinsed again twice. Thousands of carbonaceous *M. sharonae* fossils, mostly fragmentary, were isolated. Selected specimens were dehydrated to a 50% glycerine solution, then the fossil fragments were mounted on glass slides and fixed with a permanent slide mounting medium (Glycerin Gelatin). Over 600 permanent slides were created. Samples were photographed with a Nikon D90 digital camera mounted on a Leica DMRB compound microscope. High-resolution images of the mounted samples were taken using a Leica DM2500 microscope using Differential Interference Contrast, a Plan Apo 63× Oil objective, and a Nikon DS-Fi1 Digital Camera. Extended depth-of-field images were generated using Adobe Photoshop's stacking feature.

##### *Treubia lacunosa*

Fresh material of *Treubia lacunosa* (Colenso) Prosk. was collected from four localities in New Zealand's South Island. While not endangered, *T. lacunosa* is relatively rare. Because they grow in isolated colonies, collection was limited. A few individual plants were collected from several areas at each locality. Two-six 'leaf' tips per plant were used, from the 'middle' leaves only from each plant, mainly the unistratose region. Only undamaged samples were used for the analyses. They were imaged while the material was still fresh at 100x magnification using a Leica compound microscope fitted with a digital camera.

***Apotreubia nana***

Field work on *Apotreubia nana* was primarily conducted in Glacier Bay National Park under Research Permit GLBA-2023-SCI-0008. The surveys were intuitively controlled and specifically targeted two distinct habitats along the park's outermost coast. *A. nana* was found on partially shaded, liverwort-dominated, humic soil in the forested edges of low elevation peat-dominated muskegs, particularly those with a yellow cedar (*Callitropsis nootkatensis*) component. It could also be found on partially shaded, stable, and perennially damp and steeply inclined banks of incised creeks. We used LIDAR data on a GPS device in the field to help locate suitable terrain features. Samples were minimally air-dried before examination under the microscope. A "leaf" was removed from one or two plants from each collection, and a healthy and mostly unistratose apical or lateral margin was removed and mounted on a microscope slide. Photomicroscopy images were captured using a Canon R DSLR, with 5-10 images manually stacked and then combined into a single stack with Adobe Photoshop.

### Processing and microscopy of *M. polymorpha*

Tak-1 plants were grown in Jiffy-7 pots for 3-4 weeks under long-day conditions (16h/8h) at 22°C and 3800 lux. Thalli were stained with BODIPY 493/503 (Invitrogen LOT3079935) for 10 min[1,2]. A stock solution of 1 mg/mL in dimethyl sulfoxide (DMSO) was diluted to a 1 $\mu$g/mL solution in phosphate-buffered saline (PBS). Afterwards, the plants were washed for 1 min in PBS. For images obtained with both propidium iodide (PI) and BODIPY staining, plant samples were first incubated with PI (10 $\mu$g/mL) for 20 min. Afterward, the samples were washed in PBS for 1 min before being stained with BODIPY as described above. Imaging was performed using a confocal laser scanning microscope (Leica SP8) with the following settings: BODIPY (excitation: 488 nm, emission: 520-550 nm); PI (excitation: 550 nm, emission: 620-650 nm). For imaging PI and BODIPY, lasers were used in a sequential scan.

### Analysis of spatial statistics

To analyze the statistical properties of dark/OB cell patterns, we performed cell segmentation of thalli sections (Fig. S1A) with the deep learning-based cell segmentation software `cellpose`[3], using the 'cyto2' model with a diameter parameter ranging from 50 to 100 pixels (Fig. S1B). For *M. sharonae*, *T. lacunosa* and *A. nana*, segmentation was performed directly from the cell walls observed in the images. Segmentation of *M. polymorpha* was performed using cell boundaries seen through BODIPY (Fig. S7). For all species, the semi-automatic segmentation was followed by the manual correction of cells that were badly segmented. Finally, we classified cell types (dark/OB cells versus light/non-OB cells) with `ilastik`[4] (Fig. S1C). After segmentation and classification, we extracted the cell positions (centroids) from the labeled image. We then used NetworkX[5] to construct the cellular network, from which cell connectivity properties (cell neighbors, shortest paths between pairs of cells, etc.) can be extracted[6,7] (Fig. S1D). To compute the distances between neighboring dark/OB cells, we first defined dark/OB cell neighborhoods by constructing a Gabriel graph of the corresponding nodes (Fig. S1E). In a Gabriel graph, an edge between two nodes $i$ and $j$ exists only if the disk whose diameter is the line connecting the nodes $i$ and $j$ contains no other nodes[8]. This approach has previously been used to quantify the network structure of trichome patterns in *Arabidopsis*[9] and rhizoids in *Marchantia*[10]. Once the pair-wise connections between dark/OB cells have been established, we computed the shortest paths between pairs of neighboring dark/OB cells using the `shortest_path` function of the python package NetworkX[5] (Fig. S1F). With this, we have all the information to compute the cluster sizes, Moran's $I$, and the distribution of dark/OB cell distances. We used the same pipeline to analyze dark/OB cell patterns in all the models' simulations.

To quantify the spatial distribution of dark/OB cell patterns, we used the following metrics associated with each sample/simulation:

- **Dark/OB cell frequency:** the fraction of dark/OB cells over the total number of cells in that sample/simulation.

- **Frequency distribution of $k$-clusters $f_k$:** number of $k$-clusters of dark/OB cells divided by the total number of clusters of that sample/simulation.

- **Idioblast proportion:** proportion of dark/OB cell clusters of size $k = 1$, that is, from the total number of clusters, the percentage of those with a single dark/OB cell.

- **Moran's $I$ statistic**: a measure of spatial autocorrelation defined as:

$$I = \frac{N}{W} \frac{\sum_{i,j}^{N} w_{ij} \left(x_i - \bar{x}\right) \left(x_j - \bar{x}\right)}{\sum_{i}^{N} \left(x_i - \bar{x}\right)^2}$$

  where $N$ is the total number of cells in the sample/simulation, $x_i$ are the values associated with each cell ($x_i = 1$ for dark/OB cells, $x_i = 0$ for epidermal cells), $\bar{x} = \frac{1}{N} \sum_{i}^{N} x_i$ is the average value of $x_i$ (equivalent to the frequency of dark/OB cells), $w_{ij}$ is the matrix of distances between cells $i$ and $j$ (in cell units), and $W = \sum_{i,j}^{N} w_{ij}$ is a normalizing factor. Typically, Moran's $I$ ranges from -1 to 1. $I > 0$ indicates positive spatial autocorrelation or clustering, whereas $I < 0$ indicates negative spatial autocorrelation or dispersion. $I \approx 0$ indicates randomness, without regularity in the spatial pattern.

- **Distance frequency $D_j$:** the Gabriel graph defines a dark/OB cell neighborhood where each pair of cells is separated by $j$ epidermal cells (Fig. S1E,F). This separation is computed as the shortest path between pairs of dark/OB cells, i.e., the minimum number of epidermal cells connecting pairs of dark/OB cells (Fig. S1F). $D_j$ is defined as the frequency of these shortest paths. With this definition, dark/OB cells separated by $j = 0$ epidermal cells are adjacent and belong to the same cluster.

- **Average dark/OB cell distance $\langle\lambda\rangle$ and coefficient of variation $CV_\lambda$:** $\langle\lambda\rangle$ denotes the mean value of the distance frequency, $\langle\lambda\rangle \equiv \langle D_j\rangle = \sum_j jD_j$. If $\sigma_\lambda$ is its associated standard deviation ($\sigma_\lambda \equiv \sqrt{\sum_j (j-\langle\lambda\rangle)^2 D_j}$), the coefficient of variation is defined as $CV_\lambda = \sigma_\lambda/\langle\lambda\rangle$.

For consistency, we performed the statistical analysis of *T. lacunosa*, *A. nana*, and *M. polymorpha* using sections with a similar number of cells as in *M. sharonae*.

#### Model simulation and analysis

We simulated all models in irregular cell lattices that mimic the tissue layouts of mature thalli. These lattices were generated by considering a hexagonal array of cells subject to spatial noise. To do this, we created two-dimensional points arranged in a perfect triangular lattice with positions $(x,y) = (i + j/2, \frac{\sqrt{3}}{2}j)$ where $i, j$ are integers. We then randomly displaced each point a certain distance from its original location, i.e., $(x',y') = (i + j/2 + \nu\xi_x, \frac{\sqrt{3}}{2}j + \nu\xi_y)$, where $(\xi_x, \xi_y)$ are random numbers taken from a uniform distribution between 0 and 1, and $\nu$ sets the variability from the perfect hexagonal layout. If $\nu$ is small, the lattice is more or less hexagonal, with mostly six-sided cells, whereas for large values, four and three-sided cells can appear. After choosing a value for the irregularity that mimics the heterogeneity of cellular tissues ($\nu = 0.25$), a Voronoi tessellation was created from the resulting points, generating the cellular layouts used for our simulations.

#### Simulation of random patterns

To generate random patterns of dark/OB cells, every cell in a tissue of size $N_t = N_x \times N_y$ performs a binary decision to become a dark/OB cell with probability $p$ or an epidermal cell with probability $q = 1 - p$. The spatial statistics in this random model are completely determined by the parameters $p$ and $N_t$. In particular, the total number of dark/OB cells is a random variable $X$ that follows a Binomial distribution with mean $\langle X\rangle = N_t p$ and standard deviation $\sigma_X = \sqrt{N_t p(1-p)}$. The average frequency of dark/OB cells is $\langle\rho\rangle \equiv \frac{1}{N_t}\langle X\rangle = p$.

#### Model with local activation and lateral inhibition

We modeled dark/OB cell patterning using a discrete reaction-diffusion system with local activation and lateral inhibition, allowing for cell-autonomous bistability, and with diffusion restricted to the inhibitor. The model is similar to one used to describe rhizoid precursor patterning in *M. polymorpha*[10]. Each cell in our model is labeled with an index $i \in [1, \ldots, N_t]$ and is characterized by its concentrations of $u_i$ (activator) and $v_i$ (inhibitor), which dynamically change over time according to the following processes. We assume the activator has a basal synthesis rate $\alpha_0$ and can self-activate with maximum rate $\alpha$, activation threshold $k_u$, and cooperativity exponent $n_u$. Degradation of $u$ occurs both linearly at rate $d_u$ and via $v$ at rate $k_c$. The inhibitor $v$ is produced at a basal rate $\gamma_0$ and is activated by $u$ with a maximum rate $\gamma$, an activation threshold $k_v$, and a cooperativity exponent $n_v$. Finally, $v$ can be linearly degraded at rate $d_v$ and can move from cell to cell with a diffusion coefficient $D_v$. Taken together, these processes are modeled through the following set of coupled ordinary differential equations for the concentrations in each cell:

$$\frac{du_i}{dt} = \alpha_0 + \alpha \frac{u_i^{n_u}}{k_u^{n_u} + u_i^{n_u}} - k_c u_i v_i - d_u u_i \tag{1}$$

$$\frac{dv_i}{dt} = \gamma_0 + \gamma \frac{u_i^{n_v}}{k_v^{n_v} + u_i^{n_v}} - d_v v_i + D_v \Delta v_i \tag{2}$$

where $\Delta$ is a discrete Laplacian operator for an irregular lattice, defined as:

$$\Delta v_i = \frac{1}{A_i} \sum_{j \in nn_i} \frac{L_{ij}}{\gamma_{ij}} (v_j - v_i) \tag{3}$$

where $A_i$ is the area of cell $i$, $L_{ij}$ are the lengths of the shared edges between cells $i$ and $j$, $\gamma_{ij}$ the distance between centroids of cells $i$ and $j$, and $nn_i$ the nearest neighbors of cell $i$[11]. In the model, lateral inhibition is driven by the diffusive transport of $v$ to adjacent cells and the subsequent degradation of $u$ molecules. To simulate the patterns, we solved the system of coupled differential equations using the Runge-Kutta 4 method with a time step $dt = 0.02$ until the patterns became stationary ($T_{final} = 1000$; $T_{steps} = T_{final}/dt = 50000$). Boundary conditions are such that cells in the tissue boundaries only interact with their adjacent neighbors. In Table S1, we show the parameter values used for the simulations. We defined dark/OB cells as those with steady-state levels of $u$ equal to or higher than the threshold value $u_T = 1$. We used $\gamma$ and $\alpha_0$ as control parameters to analyze the stability and variability of dark/OB cell patterns (Fig. 3C; Fig. S4).

### Initial conditions

To initialize our model simulations, we considered uniformly distributed random numbers of the form $(u,v)_{t=0} = (u_0, v_0) \times \mathscr{U}[0,1]_{N_t}$, where $N_t$ denotes the number of cells in the tissue. We chose $u_0 = 1.35$ and $v_0 = 0.4$ for all our simulations.

### Parameter exploration

To identify the parameters that reproduce the spatial statistics of dark/OB cell patterns, we performed simulations on cellular tissues of size $25 \times 20$ and sampled a $100 \times 100$ grid of $(\gamma, \alpha_0)$ values, with $\gamma \in [0, 18]$ and $\alpha_0 \in [0, 1.25]$, giving 10000 parameter combinations. The remaining parameters were fixed (Table S1). For each category, we identified the pairs $(\gamma, \alpha_0)$ that generated patterns in which all metrics (dark/OB cell frequency, idioblast proportion, Moran's I, $\langle \lambda \rangle$, $CV_\lambda$) simultaneously lay within the experimental ranges. These pairs are defined by the intersection (denoted by the symbol $\bigcap$) of admissible regions for each metric, i.e., the set $S = \bigcap_l \left\{ (\gamma, \alpha_0) \mid x_l^{\min} \le X_l(\gamma, \alpha_0) \le x_l^{\max} \right\}$, where $X_l(\gamma, \alpha_0)$ denotes the value of metric $l$, and $x_l^{\min}$, $x_l^{\max}$ represent, respectively, the minimum and maximum values of each experimental metric (Figs. S5 and S6). This set constitutes a 5-dimensional hypercube that encloses all sample points. Since strictly adhering to the bounds $x_l^{\min}$, $x_l^{\max}$ may result in a small number of simulations that align with the experimental patterns, in some cases, we introduce small tolerance parameters $\delta_-$ and $\delta_+$, associated to $x_l^{\min}$, $x_l^{\max}$, respectively. This adjustment expands the acceptable range, allowing it to span from $x_l^{\min} - \delta_-$ to $x_l^{\max} + \delta_+$. In Figs. S5 and S6, we show these bounds for each metric as gray regions. In Table S2, we show $x_l^{\min}$, $x_l^{\max}$, $\delta_-$, and $\delta_+$ associated with each metric for each pattern type in *M. sharonae*, and for the extant taxa.

#### Bifurcations and linear stability analysis

We performed linear stability analysis[12,13] of our model to find the parameter space regions where patterns emerge spontaneously from a homogeneous state due to small perturbations (Fig. 3C; region 6). For this analysis, we considered a hexagonal lattice of size $25 \times 20$, where the spatial position of each cell is defined by the indices $j,k$. To find the stability of a homogeneous state, we first numerically compute the stationary homogeneous states by computing the fixed points of the uncoupled system. We denote a stationary homogeneous solution as the vector $\mathbf{x}_h \equiv (u_h, v_h)$. Next, we consider a perturbation of the form $\delta\mathbf{x}_{jk} = \mathbf{x}_{jk} - \mathbf{x}_h$, where $\mathbf{x}_{jk} \equiv (u_{jk}, v_{jk})$ is a vector of all the system's variables. We assume perturbations of the form:

$$\delta\mathbf{x}_{jk} = \sum_p^{N_x}\sum_q^{N_y} \mathbf{A}_{pq} e^{2\pi \mathrm{i}\left(\frac{jp}{N_x}+\frac{kq}{N_y}\right)} e^{\sigma_{pq}t} \tag{4}$$

where $p,q$ denote the Fourier modes, i the imaginary number, $\mathbf{A}_{pq}$ is the modes' amplitude, and $\sigma_{pq}$ is their growth rate. The dynamics of these perturbations are analyzed for the linearized version of the model around the homogeneous state. Therefore, the growth rates $\sigma_{pq}$ correspond to the eigenvalues of the Jacobian matrix:

$$\mathbb{J}_{pq} = \begin{pmatrix} \alpha n_u \frac{k_u^{nu} u_h^{nu-1}}{(k_u^{nu}+u_h^{nu})^2} - k_c v_h - d_u & -k_c v_h \\ \gamma n_v \frac{k_v^{nv} u_h^{nv-1}}{(k_v^{nv}+u_h^{nv})^2} & -d_v - 4D_v(1-\Omega_{pq}) \end{pmatrix} \tag{5}$$

where $\Omega_{pq} \equiv \frac{1}{3}\left[\cos\left(\frac{2\pi p}{N_x} - \frac{2\pi q}{N_y}\right) + \cos\left(\frac{2\pi p}{N_x}\right) + \cos\left(\frac{2\pi q}{N_y}\right)\right]$. This quantity ranges from $\Omega_{pq} \in [-\frac{1}{2}, 1]$, having its minimum when all terms in the brackets equal $-\frac{1}{2}$, and its maximum when they equal to 1. The corresponding eigenvalue equation takes the form of a rank-2 polynomial $\lambda^2 + A\lambda + B = 0$, where the coefficients $A,B$ are constants associated with the Jacobian $\mathbb{J}_{pq}$ evaluated at the homogeneous states. The homogeneous state is linearly unstable when the real part of any of the growth rates $\sigma_{pq}$ is positive ($\mathrm{Re}(\sigma_{pq}) > 0$). We solved the eigenvalue equation numerically for different values of the parameters $(\gamma, \alpha_0)$ to find for which parameter values the condition of instability ($Re(\sigma_{pq}) > 0$) is met for any $(p,q)$. The spatial modes that first become unstable (those with $\Omega_{pq} = -\frac{1}{2}$) are $p = N_x/3$ and $q = 2N_y/3$ (and their reciprocals $p = 2N_x/3$ and $q = N_y/3$), that is, salt-and-pepper patterns. In Fig. S4B, we depict the Turing bifurcations (Re($\lambda$)= 0 for inhomogeneous modes, Re($\lambda$)< 0 for homogeneous modes; magenta curves) and the regions of spontaneous pattern formation (Re($\lambda$)> 0 for inhomogeneous modes, Re($\lambda$)< 0 for homogeneous modes; pink and magenta regions) in the plane $(\gamma, \alpha_0)$. To compute the bifurcations in Fig. 3C and Fig. S4, we used custom-made software and the Julia package BifurcationKit[14].

#### Patterning regimes of the system

In Fig. 3C, we show all the possible patterning regimes in the $(\gamma, \alpha_0)$ plane. If $\alpha_0$ (production of the activator) is small, dark cells do not appear (region 1). Conversely, if $\alpha_0$ is large, all cells become dark. For intermediate values of $\alpha_0$ and $\gamma$, patterns can occur. In Fig. S4, we show all these patterning regimes and their associated bifurcations and instabilities. Without diffusion ($D_v = 0$), the system is equivalent to a single-cell system and exhibits a region of cell-autonomous bistability, bounded by saddle-node bifurcations (Fig. S4A; gray region). These states correspond to homogeneous states in the spatially extended system. In the bistable region, both homogeneous states (no dark cells and all dark cells) are linearly stable, and the emergence of dark cell patterns depends exclusively on the initial conditions. When diffusion is introduced, diverse patterning regimes emerge (Fig. S4B). Linear stability analysis reveals a region of spontaneous pattern formation enclosed by subcritical Turing bifurcations (Fig. S4B; pink region). Here, patterns emerge spontaneously, span the whole tissue, and tend to display spatial regularity (Fig. S4D). A small part of this region coexists with the low-$u$ homogeneous stable state (magenta region). Here, high-$u$ states can lose stability due to small inhomogeneous perturbations, whereas the low-$u$ homogeneous state is linearly stable. In the gray region, both homogeneous states (low-$u$ and high-$u$) are linearly stable, and coincides with the same region in the cell-autonomous case. Outside regions of spontaneous pattern formation, regular patterns can still emerge (localized states I, II; turquoise regions). In these regions, pattern solutions coexist with homogeneous states, are nonlinear, and depend on initial conditions. Localized patterning regions are enclosed by saddle-node bifurcations (dashed turquoise curves) corresponding to the pattern states that first become stable, that is, the state with all dark cells and a single light cell (localized patterns I), and the state with a single dark cell in a tissue of light cells (localized patterns II).

### Supplementary tables

| Parameter | Description | Value (a.u.) |
|---|---|---|
| $\alpha_0$ | basal transcription rate of $u$ | variable |
| $\alpha$ | maximum value of $u$ self-activation | 2 |
| $k_u$ | threshold of $u$ self-activation | 1.2 |
| $n_u$ | cooperativity exponent of $u$ self-activation | 4 |
| $k_c$ | repression rate of $u$ by $v$ | 1 |
| $d_u$ | $u$ degradation | 1 |
| $\gamma_0$ | basal transcription rate of $v$ | 0.1 |
| $\gamma$ | maximum value of $v$ activation by $u$ | variable |
| $k_v$ | threshold of $v$ activation by $u$ | 2 |
| $n_v$ | cooperativity exponent of $v$ activation by $u$ | 4 |
| $d_v$ | $v$ degradation | 1 |
| $D_v$ | diffusion coefficient of $v$ | 4 |
| $u_T$ | threshold defining dark cell fate | 1.0 |
| $u_0$ | initial concentration of $u$ | $1.35 \times \mathscr{U}[0,1]_{N_t}$ |
| $v_0$ | initial concentration of $v$ | $0.4 \times \mathscr{U}[0,1]_{N_t}$ |

**Table S1:** Parameters used for the model with local activation and lateral inhibition, their description, and their values (in arbitrary units). The parameters used for the analysis ($\alpha_0$ and $\gamma$) are shaded in gray. $\mathscr{U}[0,1]_{N_t}$ denotes a uniform distribution of random numbers between 0 and 1 in a tissue of $N_t$ cells.

| Metric | Patterns | $x_l^{min}$ | $x_l^{max}$ | $\delta_-$ | $\delta_+$ |
|---|---|---|---|---|---|
| dark/OB cell frequency | *M. sharonae* (type I) | 0.194 | 0.263 | 0 | 0.02 |
| | *M. sharonae* (type II) | 0.076 | 0.189 | 0 | 0 |
| | *M. sharonae* (type III) | 0.148 | 0.299 | 0 | 0 |
| | *M. sharonae* (type IV) | 0.005 | 0.024 | 0 | 0 |
| | *T. lacunosa* | 0.092 | 0.228 | 0 | 0 |
| | *A. nana* | 0.097 | 0.219 | 0 | 0 |
| | *M. polymorpha* | 0.033 | 0.045 | 0 | 0 |
| idioblast proportion | *M. sharonae* (type I) | 0.82 | 0.96 | 0.05 | 0.05 |
| | *M. sharonae* (type II) | 0.70 | 1 | 0 | 0 |
| | *M. sharonae* (type III) | 0.67 | 0.86 | 0 | 0 |
| | *M. sharonae* (type IV) | 0.91 | 1 | 0 | 0 |
| | *T. lacunosa* | 0.56 | 1 | 0 | 0 |
| | *A. nana* | 0.65 | 0.89 | 0 | 0 |
| | *M. polymorpha* | 0.94 | 1 | 0.02 | 0 |
| Moran's I | *M. sharonae* (type I) | -0.205 | -0.139 | 0.05 | 0.05 |
| | *M. sharonae* (type II) | -0.176 | -0.011 | 0 | 0 |
| | *M. sharonae* (type III) | -0.140 | -0.035 | 0 | 0 |
| | *M. sharonae* (type IV) | -0.002 | 0.015 | 0.005 | 0.005 |
| | *T. lacunosa* | -0.185 | -0.015 | 0 | 0 |
| | *A. nana* | -0.136 | -0.052 | 0 | 0 |
| | *M. polymorpha* | -0.035 | -0.012 | 0.01 | 0.01 |
| average dark/OB cell distance $\langle\lambda\rangle$ | *M. sharonae* (type I) | 2.13 | 2.53 | 0.3 | 0.3 |
| | *M. sharonae* (type II) | 2.45 | 3.87 | 0 | 0 |
| | *M. sharonae* (type III) | 1.96 | 2.52 | 0 | 0 |
| | *M. sharonae* (type IV) | 5.88 | 18.00 | 0 | 0 |
| | *T. lacunosa* | 2.04 | 3.32 | 0 | 0 |
| | *A. nana* | 2.09 | 3.28 | 0 | 0 |
| | *M. polymorpha* | 4.57 | 5.69 | 0.1 | 0.1 |
| $CV_\lambda$ | *M. sharonae* (type I) | 0.25 | 0.35 | 0.05 | 0.05 |
| | *M. sharonae* (type II) | 0.26 | 0.50 | 0 | 0 |
| | *M. sharonae* (type III) | 0.34 | 0.47 | 0 | 0 |
| | *M. sharonae* (type IV) | 0 | 0.45 | 0 | 0 |
| | *T. lacunosa* | 0.27 | 0.48 | 0 | 0 |
| | *A. nana* | 0.31 | 0.44 | 0 | 0 |
| | *M. polymorpha* | 0.28 | 0.0.44 | 0.02 | 0.02 |

**Table S2:** Table showing the minimum ($x_l^{min}$), maximum ($x_l^{max}$), and tolerance values ($\delta_-$, $\delta_+$) for each metric and pattern type in *M. sharonae*, and extant taxa.

### Supplementary figures

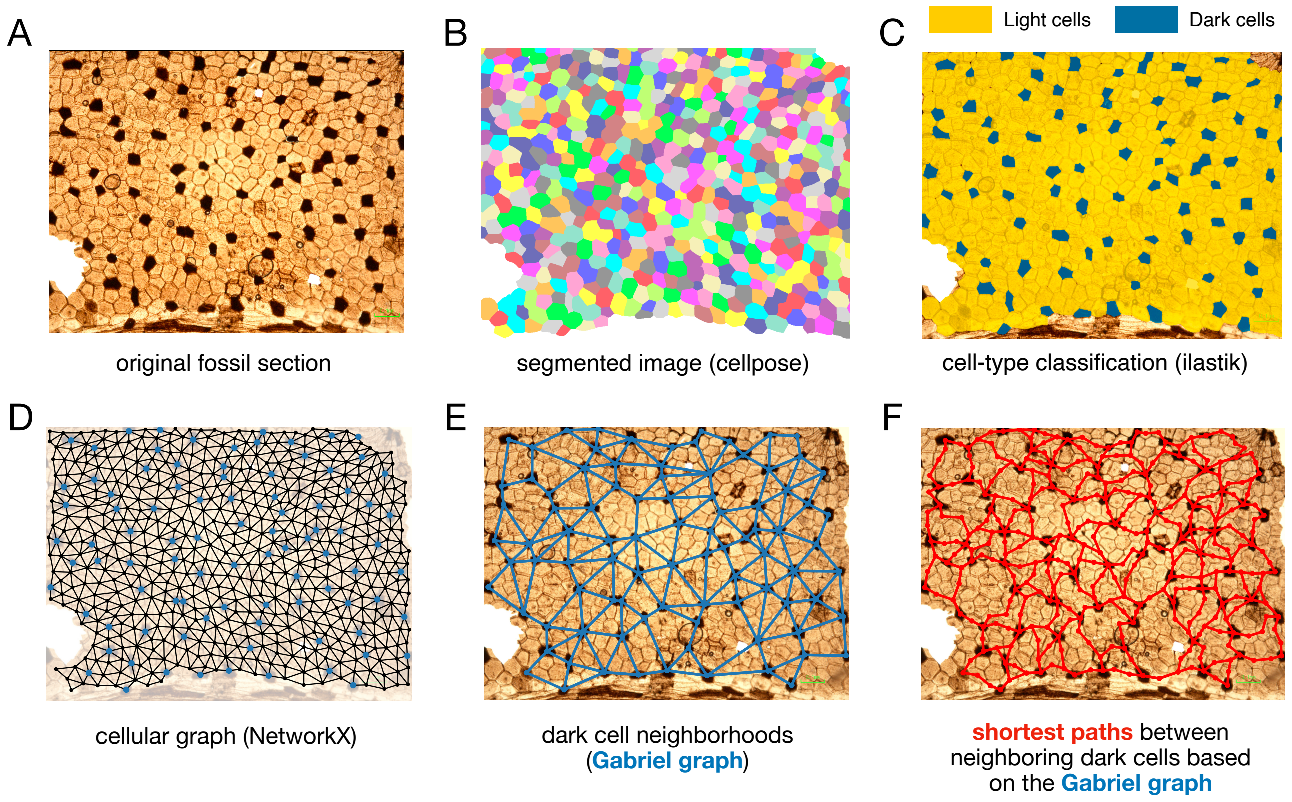


**Figure S1: Pipeline for quantifying dark/OB cell patterns.** **A.** Example fossil section used for segmentation, annotation, and analysis. **B.** Cellular segmentation with `cellpose`[3], showing all segmented cells in different colors. **C.** Classification of cell types (dark and light) using `ilastik`[4]. **D–F.** Pipeline for spatial analysis of segmented tissues. First, we used NetworkX[5] to construct a cellular graph with edges connecting the centroids of adjacent cells (D). Then, we constructed a Gabriel graph determining dark cell neighborhoods (E). Dark cell distances can be computed through the shortest paths between pairs of dark cells (F). The same pipeline was used for analyzing dark/OB cell patterns in the simulations.

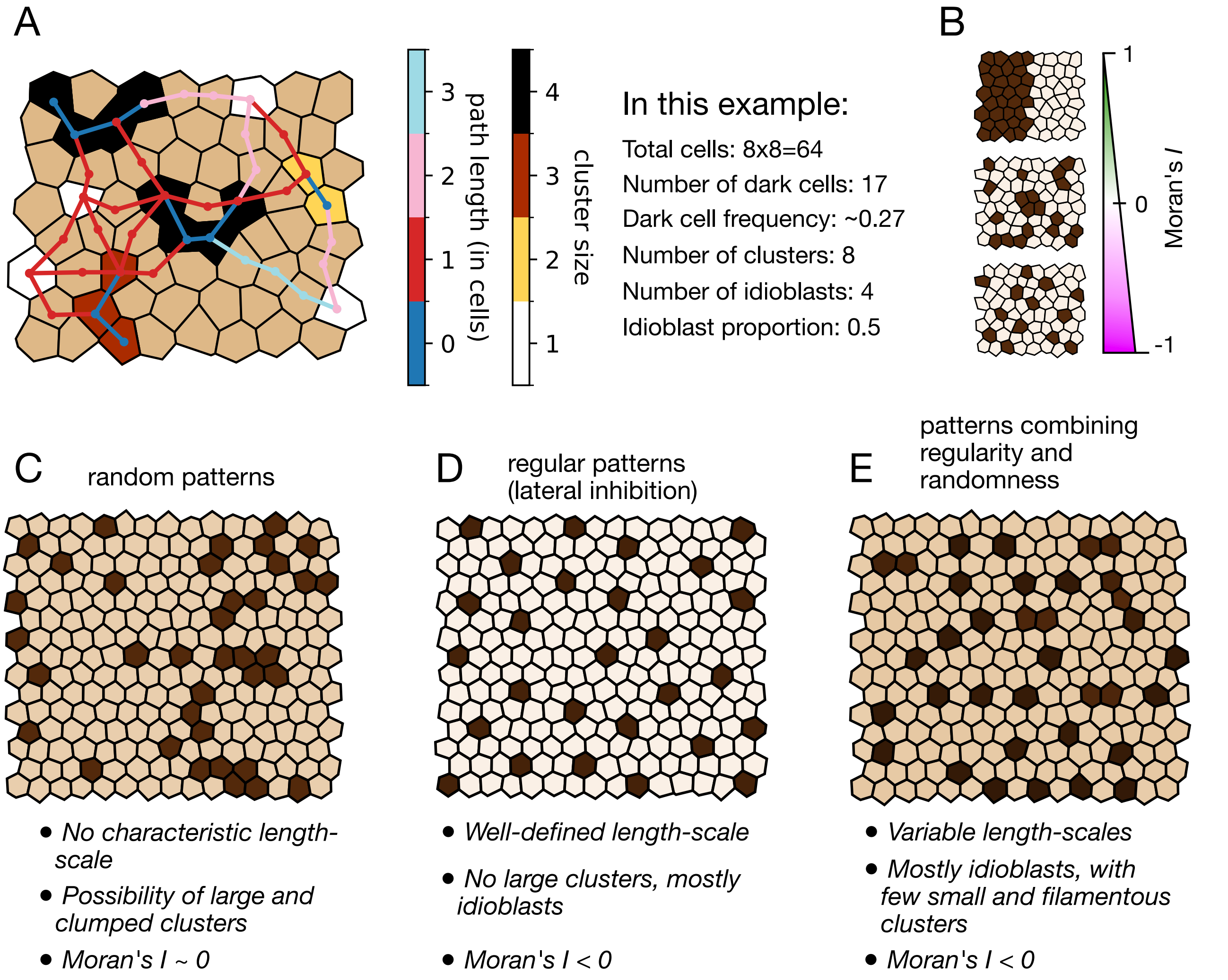


**Figure S2: Statistical metrics and their meaning for different pattern types. A.** Example tissue with several dark/OB cell clusters, showcasing cluster sizes in different colors and the shortest paths between pairs of neighboring dark cells. **B.** Intuitive sketch showing how Moran's $I$ changes depending on the pattern. Completely clustered patterns have Moran's $I \approx 1$; random patterns have Moran's $I \approx 0$; regular patterns have Moran's $I \approx -1$. **C–E** Each pattern type has a statistical signature that can be used to infer the mechanisms underlying its patterning. Random patterns have no characteristic length scale, and Moran's $I \approx 0$ (C); regular patterns are associated with lateral inhibition, with a well-defined length scale, no large clusters, and negative Moran's $I$ (D); mixed patterns are associated with a combination of randomness and lateral inhibition, with variable length scales, filamentous clusters, and negative Moran's $I$.

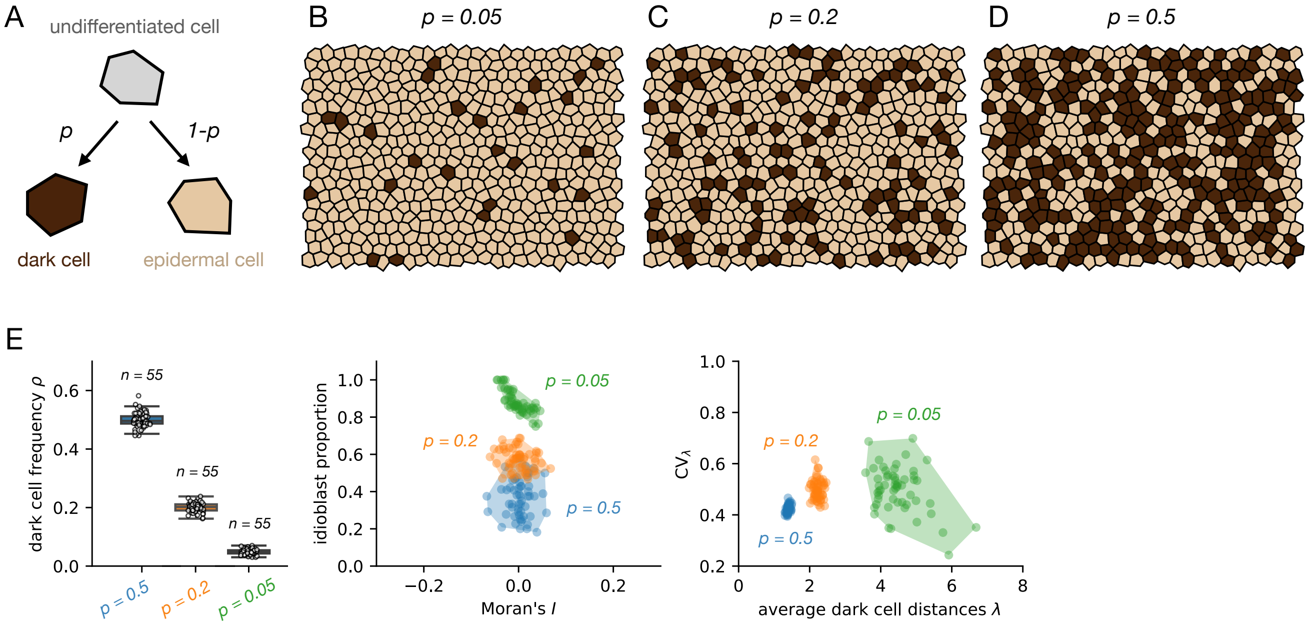


**Figure S3: Random models and their statistics.** **A.** In a random model of cell specification, every undifferentiated cell (light gray) can differentiate into a dark cell or an epidermal cell with probability $p$ and $1-p$, respectively. **B–D.** Typical dark cell patterns obtained with the random model, for $p = 0.05$ (B), $p = 0.2$ (C), and $p = 0.5$ (D). **E.** Spatial statistics (dark cell frequency, Moran's $I$, idioblast proportion, average dark cell distance $\langle\lambda\rangle$ and $CV_\lambda$) of $n = 55$ patterns for each value of $p$ detailed above. Regardless of the value $p$, random patterns have a Moran's $I$ around 0, and their idioblast proportion is inversely proportional to $p$. The lower the $p$, the larger the distance between pairs of dark cells, whereas $CV_\lambda$ remains approximately constant.

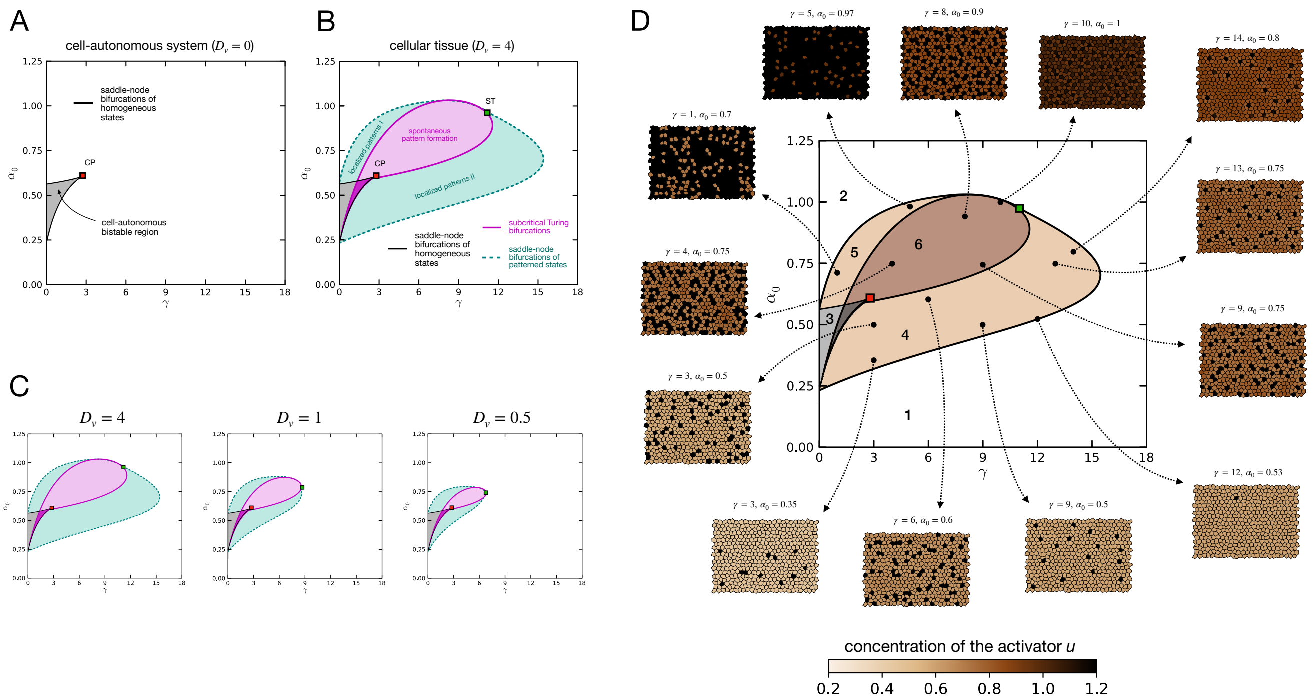


**Figure S4: Bifurcation diagrams and patterning regimes of the mathematical model. A.** Stability diagram of the mathematical model for a single cell, showing a region of cell-autonomous bistability enclosed by saddle-node bifurcations that collide in a cusp point (CP). **B.** Stability diagram of the mathematical model for a cellular tissue, showing each patterning region and the bifurcations separating them. Subcritical Turing bifurcations enclose the region of spontaneous pattern formation (pink curve). Overlapping with the region of cell-autonomous bistability, there is a small region (magenta region) where the high-$u$ state can lose stability due to inhomogeneous perturbations. In this region, patterned states coexist with the low-$u$ homogeneous state. The two regions of localized patterns are delimited by saddle-node bifurcations (dashed turquoise curves) associated with the states with a single light cell (localized patterns I) and the state with a 1 single dark cell (localized patterns II). The red square denotes a cusp point (CP) where the two saddle-node bifurcations associated with the homogeneous states collide. The green square denotes the saddle-Turing (ST) point, where the two saddle-node bifurcations associated with the 1-cell states collide with the subcritical Turing bifurcation. **C.** The effect of different diffusion coefficients ($D_v = 4, 1, 0.5$) on patterning regions. As diffusion becomes smaller, the patterning regions decrease in size. **D.** Stationary dark cell patterns obtained in different parameter regions. In the region of spontaneous pattern formation, patterns exhibit spatial regularity, but their density can vary. In regions of localized patterns, multiple patterning states can coexist with the homogeneous state, enabling the formation of patterns that are restricted to specific areas rather than spanning the whole tissue. The remaining parameters and initial conditions for obtaining these patterns are detailed in Table S1.

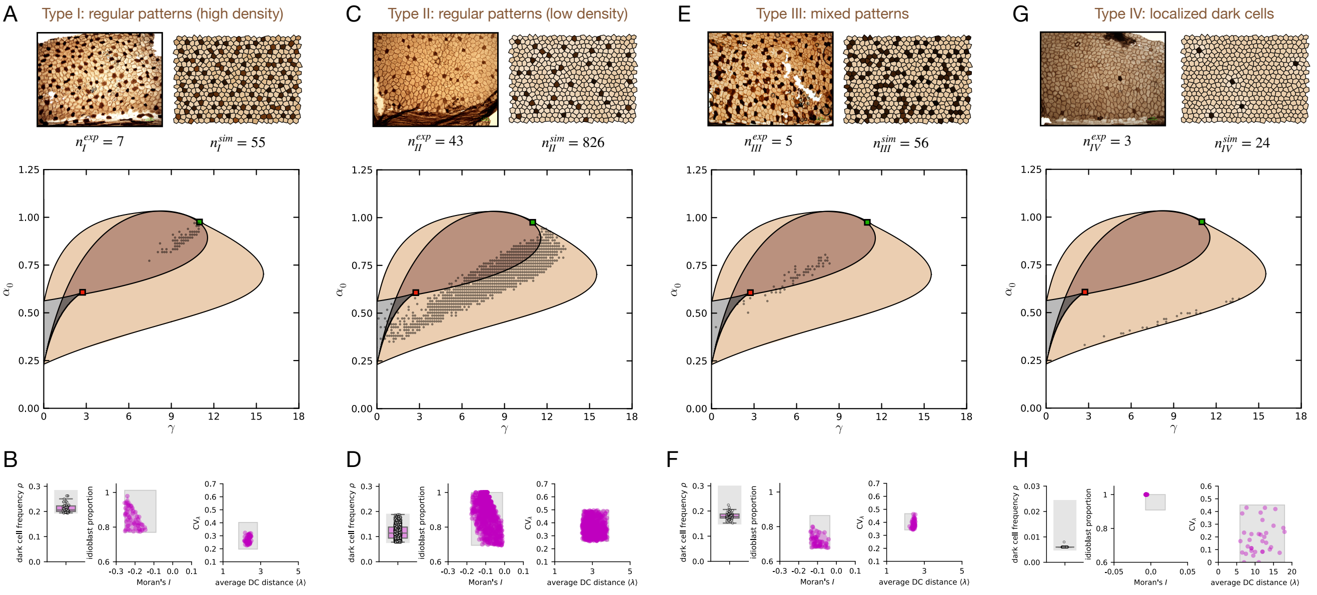


**Figure S5: Pattern types in *M. sharonae*, their associated model parameter values, and their statistical analysis. A–B.** Type I: regular, high dark cell density patterns appear in the region of spontaneous pattern formation. Due to this regularity, Moran's $I$ is negative, and the average dark cell distance is very consistent, with small $CV_\lambda$. **C–D.** Type II: Regular, low dark cell density patterns appear between the regions of spontaneous pattern formation and the region of localized patterns. These patterns are less regular than Type I, with Moran's $I$ closer to 0 and larger $CV_\lambda$. **E–F.** Type III: mixed patterns appear in the region of spontaneous pattern formation close to the bistability region. **G–H.** Type IV: patterns with only a few dark cells appear only in the region of low-density localized patterns close to the bifurcation boundary. Stability diagrams are as in Fig. 3C. Black scattered points mark the values of $(\gamma, \alpha_0)$ reproducing the dark cell patterns of each type. Gray boxes in B,D,F,G mark the regions between $x_l^{\min} - \delta_-$ and $x_l^{\max} + \delta_+$ for each metric (see Table S2).

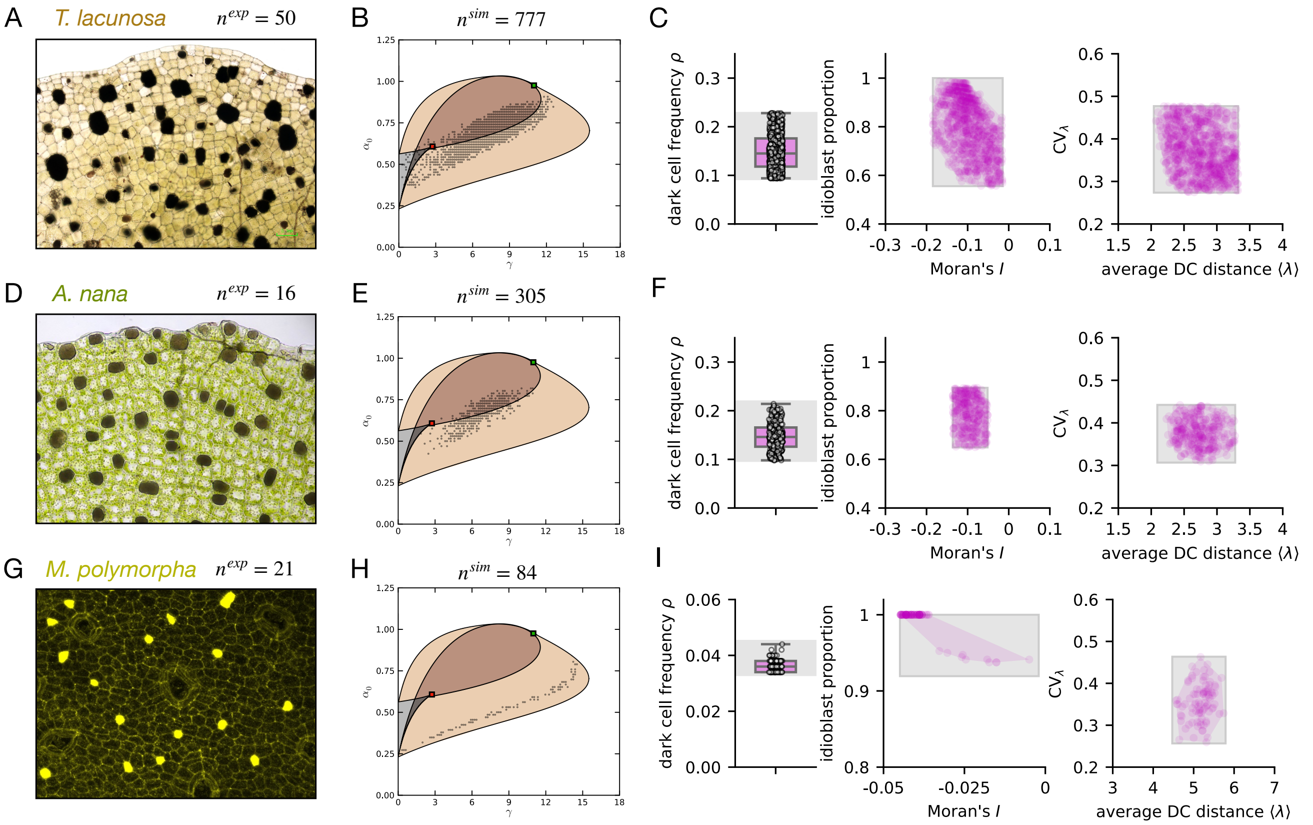


**Figure S6: Values of $(\gamma, \alpha_0)$ associated with OB cell patterns in extant liverworts. A–C.** In *T. lacunosa*, OB cell patterns appear in the regions of spontaneous and localized patterns. **D–F.** Patterns in *A. nana* appear in similar regions than *T. lacunosa*, highlighting their phylogenetic similarity. **G–I.** In *M. polymorpha*, OB cells appear with a very low frequency, in the region of localized patterns close to the bifurcation boundary. Black scattered points mark the values of $(\gamma, \alpha_0)$ reproducing OB cell patterns in each species. Gray boxes in C,F,I mark the regions between $x_l^{\min} - \delta_-$ and $x_l^{\max} + \delta_+$ for each metric (see Table S2).

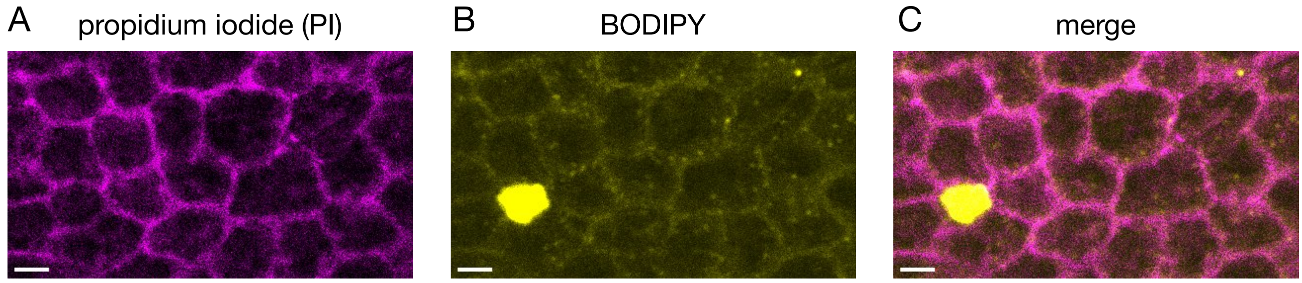


**Figure S7: BODIPY and PI staining in the lobe region of a *Marchantia* thallus. A.** Confocal image showing cell walls in the PI channel. **B.** Cell boundaries seen through BODIPY staining. **C.** Merge, confirming that BODIPY enables the detection of cell boundaries in the observed region. Scale bars = 1 mm.

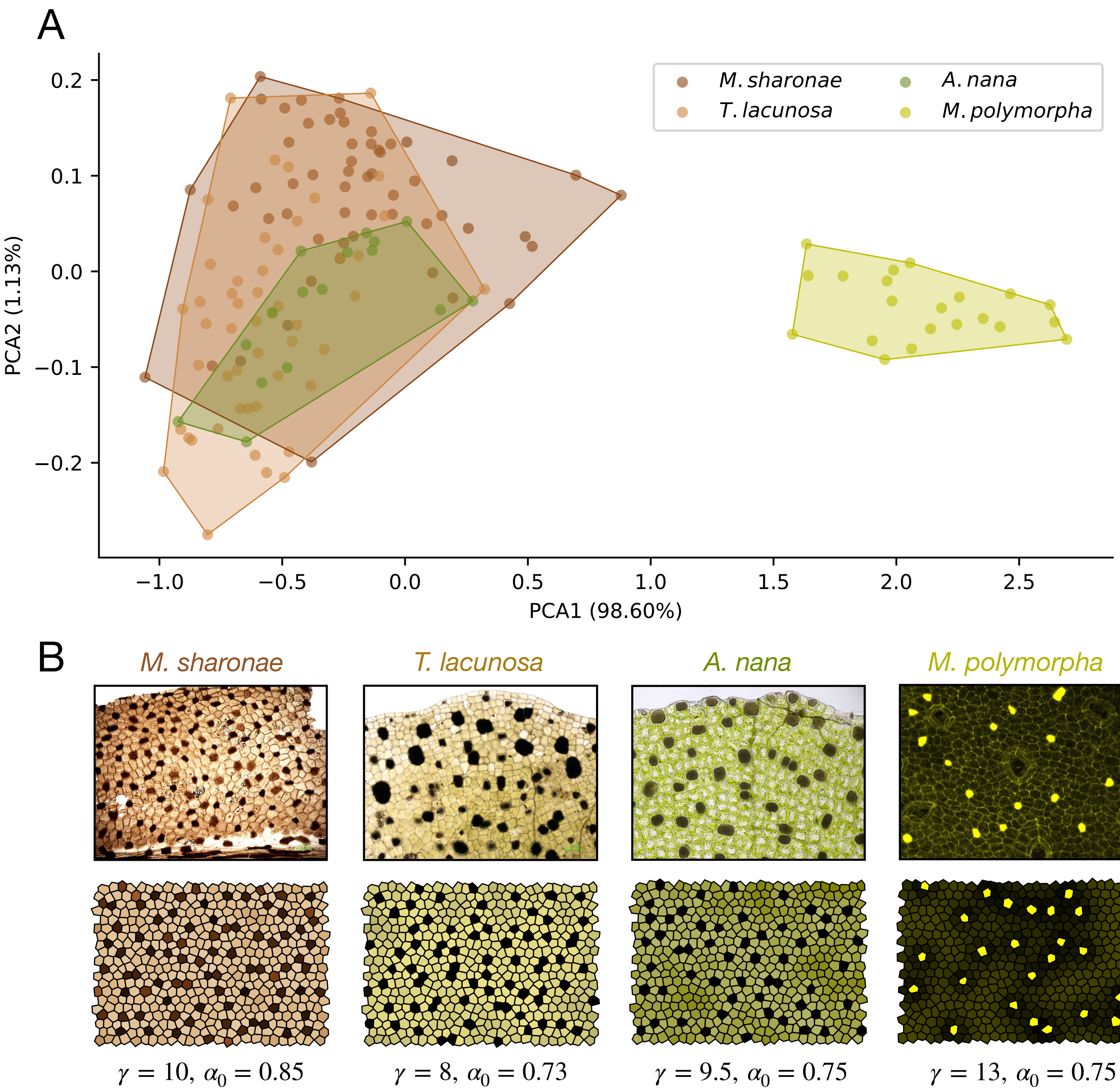


**Figure S8: OB cell distribution in different species.** **A.** Principal component analysis (PCA) showing how different species cluster in the space of (PCA1, PCA2). PCA was performed on the five quantification measures of the dark cells/OB patterns using the `scikit-learn` package in python[15]. Patterns in *M. sharonae*, *T. lacunosa* and *A. nana* overlap in this space, showcasing their similarity. OB cell patterns in *M. polymorpha* appear far from the rest due to their significantly lower OB cell frequency. **B.** Patterns (top) and associated example simulations (bottom) reproducing the spatial statistics of each species, together with the parameters $(\gamma, \alpha_0)$ used. The rest of the parameters and initial conditions for the simulations are detailed in Table S1.

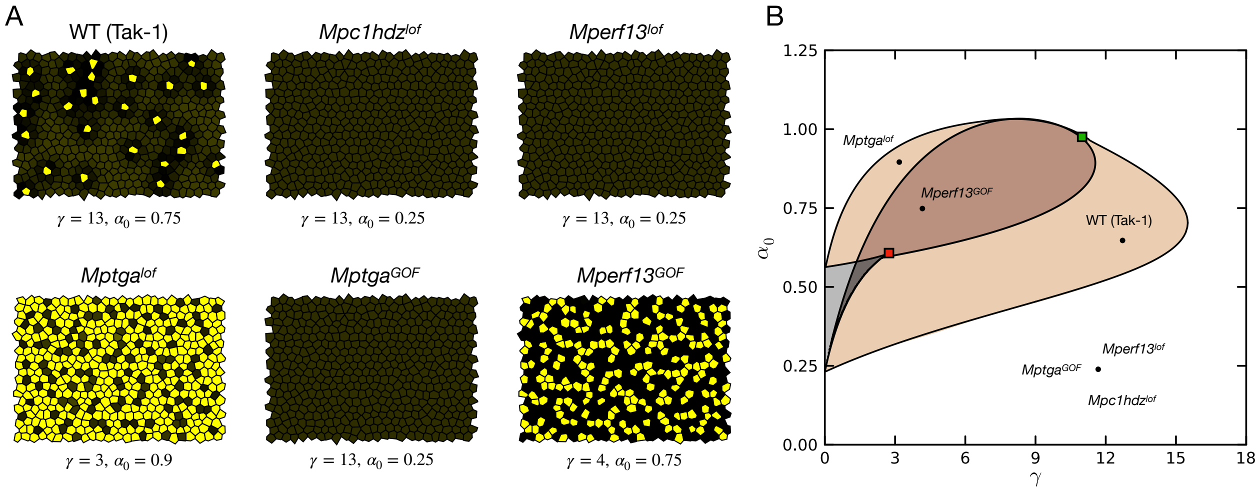


**Figure S9: The model can reproduce OB cell patterns in gain and loss-of-function genes controlling OB cell specification. A.** Different patterns obtained with the model showing the qualitative OB cell distribution in WT and different mutants. In *Mpc1hdz*$^{lof}$ and *Mperf13*$^{lof}$, OBs do not appear due to the lack of their activators. In *Mptga*$^{lof}$, the number of OBs increases dramatically, as TGA is an inhibitor of OB cell specification. This increase also occurs in *Mperf13*$^{GOF}$, but more weakly. **B.** Parameter values used for the simulations in A. In our framework, different mutants reside in different regions of the parameter space, therefore exhibiting different OB cell patterns. The rest of the parameters and initial conditions are detailed in Table S1.

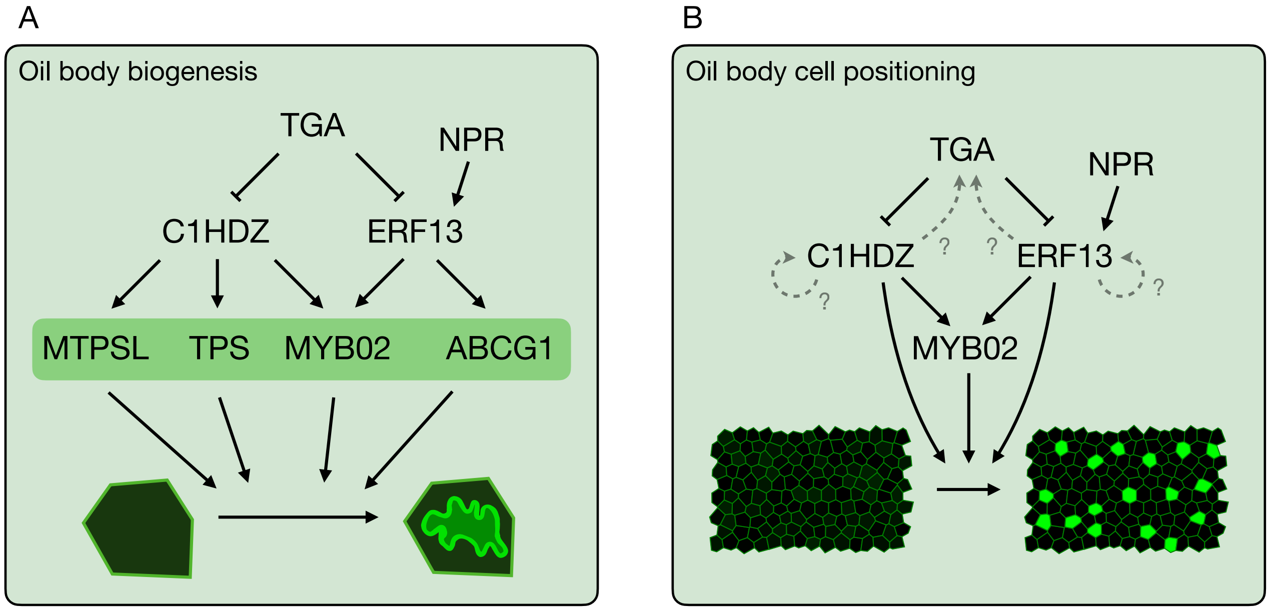


**Figure S10: Gene regulatory network controlling OB biogenesis and positioning.** **A.** Known regulatory pathway controlling OB biogenesis. **B.** Transcription factors controlling the spatial distribution of OB cells and hypothetical feedback interactions required to generate cell-type patterns. Normal arrows represent activation, and blunt arrows represent inhibition.
